# Genetic architecture underlying response to the fungal pathogen *Dothistroma septosporum* in lodgepole pine, jack pine, and their hybrids

**DOI:** 10.1101/2024.04.24.590931

**Authors:** Mengmeng Lu, Nicolas Feau, Brandon Lind, Dragana Obreht Vidakovic, Pooja Singh, Sally N. Aitken, Richard C. Hamelin, Sam Yeaman

**Affiliations:** Department of Biological Sciences, University of Calgary, 507 Campus Drive NW, Calgary, AB T2N 1N4, Canada; Department of Biological Sciences, University of Notre Dame, 100 Galvin Life Sciences, Notre Dame, IN 46556, USA; Department of Forest and Conservation Sciences, University of British Columbia, 3041-2424 Main Mall, Vancouver, BC V6T 1Z4, Canada; Canadian Forest Service, Pacific Forestry Centre, Natural Resources Canada, Victoria, BC V8Z 1M5, Canada; Aquatic Ecology & Evolution Division, Institute of Ecology and Evolution, University of Bern, Bern CH-3012, Switzerland; Center for Ecology, Evolution & Biogeochemistry, Swiss Federal Institute of Aquatic Science and Technology (EAWAG), Kastanienbaum CH-6047, Switzerland; Institut de Biologie Intégrative et des Systèmes, Université Laval, Pavillon Charles-Eugène- Marchand 1030, avenue de la Médecine, Québec City, Québec G1V 0A6, Canada

**Keywords:** Conifer, fungal pathogen resistance, *F*_ST_, hybrid, introgression, pool-GWAS

## Abstract

In recent decades, *Dothistroma* needle blight (DNB), a pine tree disease caused by the fungal pathogen *Dothistroma septosporum*, has severely damaged lodgepole pine (*Pinus contorta* Dougl. ex. Loud.) in British Columbia, Canada, and raised health concerns for jack pine (*Pinus banksiana* Lamb.). The pathogen has already shown signs of host shift eastward to the hybrid populations between lodgepole pine and jack pine (*Pinus contorta* × *P. banksiana*), and possibly into pure jack pine. However, we have little knowledge about mechanisms of resistance to *D. septosporum*, especially the underlying genetic basis of variation in pines. In this study, we conducted controlled inoculations to induce infection by *D. septosporum* and performed a genome-wide case-control association study with pooled sequencing (pool-seq) data to dissect the genetic architecture underlying response in lodgepole pine, jack pine, and their hybrids. We identified candidate genes associated with *D. septosporum* response in lodgepole pine and in hybrid samples. We also assessed genetic structure in hybrid populations and inferred how introgression may affect the distribution of genetic variation involved in *D. septosporum* response in the studied samples. These results can be used to develop genomic tools to evaluate DNB risk, guide forest management strategies, and potentially select for resistant genotypes.

## 1 INTRODUCTION

Pests and pathogens constantly threaten forest trees, and their impacts are changing as climates are altered (Seidl et al., 2017; Simler-Williamson et al., 2019). Knowledge about the genetic architecture of pathogen resistance and the identified candidate resistance (R) genes provides the basis for genomic breeding to improve tree resilience to forest pathogens. Genes and alleles that confer disease resistance or tolerance can eventually be used to develop genomic tools for rapidly selecting resistant genotypes (Isabel et al., 2019). Current breeding practices for forest trees involve recurrent selection, which includes repeated cycles of selection, breeding, and testing, to search for genotypes with better disease resistance, and then deployment of these selected tree genotypes (White, 2004). This takes years of effort to design and implement.

Increasingly, genome editing is used to develop crops resistant to disease (van Esse et al., 2020). For fruit trees, CRISPR/Cas9 system has been applied to enhance the disease resistance in citrus (Jia et al., 2017; Wang et al., 2019), apple (Zhou et al., 2020), and others (Min et al., 2022).

Given the long generation time in conifers and the frequent threats from diseases and insects, identifying R genes that can be targeted for selection or editing is a useful step in developing resistant genotypes for reforestation.

*Dothistroma* needle blight (DNB), which is caused by the fungal pathogen *Dothistroma septosporum*, is a foliar disease of a wide range of pine trees (Gibson, 1972), with infected trees showing reddish-brown bands in needles, defoliation, and growth reduction. DNB has caused increased damage to lodgepole pine (*Pinus contorta* Dougl. ex. Loud., LP) in recent decades in Western Canada (Dale et al., 2011; Woods, 2003), and is now expanding its range eastwards, threatening jack pine (*Pinus banksiana* Lamb., JP). In severe cases, DNB causes extensive mortality, even amongst mature trees in LP plantations (Woods, 2003). Historically, DNB had only minor impacts on native forest trees in North America, but emerged as a severe forest disease during 1950s to 1960s in *Pinus radiata* plantations in Africa, New Zealand, and South America (Gibson, 1972). Recently, DNB has caused increased outbreak incidence and host range expansion in North America and Europe, which is likely correlated with climate change (Boroń et al., 2021; Welsh et al., 2009; Woods et al., 2005). In North America, while *D. septosporum* tends to be more common in moist western forests closer to the Pacific Ocean, *Dothistroma*-like symptoms were observed in drier forests of LP and natural hybrids between LP and JP (*Pinus contorta* × *P. banksiana*, LP × JP) in northern Alberta in 2012 and 2013 with further tests confirming *D. septosporum* was the cause of the disease observed (Ramsfield et al., 2021). Feau et al. (2021) performed a controlled inoculation experiment and demonstrated that LP, JP, and LP × JP are susceptible to *D. septosporum*, with JP showing a higher disease severity than LP or LP × JP, though there is no report showing DNB infection in natural JP forests. Considering *D. septosporum*’s eastward shift reaching towards the natural range of JP and JP’s high susceptibility to *D. septosporum*, it is important to understand how these species respond to DNB infection.

The evolutionary history and biogeography of JP and LP have likely played an important role in shaping the genetic basis of resistance to DNB. JP is widely distributed across boreal forests in eastern North America, extending in Canada from Northwest Territories to Nova Scotia (Rudolph & Laidly, 1990), areas largely outside of the historical range of *Dothistroma*. LP and JP evolved in allopatry, with an estimated divergence time of ∼8 million years before present (Hao et al., 2015), but hybridized in areas of contact, primarily in central and northwestern Alberta (Burns et al., 2019; Rudolph & Yeatman, 1982). Following secondary contact of these two species, additional patchy regions of introgression have been developed in central Alberta north through to the Northwest Territories and northeastern British Columbia to the Alberta-Saskatchewan border (Burns et al., 2019). Natural LP × JP hybrids display a wide range of phenotypes that are intermediate or more typical of one parent, depending on the level of introgression (Wood et al., 2009; Yeatman & Teich, 1969).

In light of the geographic distribution of LP and JP, the longer history of host-pathogen coevolution in LP might explain its stronger resistance to DNB infection, compared to JP. A former study has evidenced coevolution between *Cronartium harknessii* lineages with LP and JP, with genetic basis underlying both pathogen virulence and host resistance (McAllister et al., 2022). For DNB infection caused by *D. septosporum*, though we lack direct evidence to prove a coevolutionary history between *D. septosporum* and different pine hosts, we have already identified candidate resistance genes to *D. septosporum* by RNA-seq analysis on experimentally infected LP seedlings (Lu et al., 2021). These candidate genes include 43 genes that are present in the plant-pathogen interaction pathway, as well as nine R genes that contained sites under positive selection. It seems that both constitutive (baseline) and induced (defenses activated after being attacked) defenses have been developed in LP. However, little is known about genetic basis of resistance to *D. septosporum* in JP. When subjected to other biotic agents, LP and JP showed unique monoterpene profiles in response to mountain pine beetle (Hall et al., 2013) and significant differences in chitinase gene expression in response to the fungal pathogens *Grosmannia clavigera* and *Cronartium harknessii* (Peery et al., 2021). These previous studies revealed the species-specific pathogen responses in a focused set of gene families. Nevertheless, since many complex traits are determined by many genes of small effect, to identify the causal genes underlying disease response traits, a genome-wide gene survey might give us a high-resolution perspective (Alonso-Blanco & Méndez-Vigo, 2014; Tam et al., 2019).

To dissect the genetic architecture of pathogen resistance and to identify candidate genes, researchers have employed genetic markers, mainly SNPs, and methods like quantitative trait locus (QTL) mapping and genome-wide association analysis (GWAS). Polygenic disease responses, comprising numerous loci of small effect, appear to be the rule (Stocks et al., 2019). Large numbers of SNPs were associated with pitch canker resistance in *Pinus taeda* L. (De La Torre et al., 2019; Lu et al., 2017; Quesada et al., 2010), with white pine blister rust resistance in *Pinus lambertiana* Dougl. (Weiss et al., 2020), and with *Heterobasidion* root rot resistance in *Picea abies* (L.) Karst. (Capador-Barreto et al., 2021). To identify the candidate R genes, most GWAS studies on forest trees employ individual-based methods, with both phenotypic and genotypic data collected on each sampled individual. Case-control methods have been developed to study disease where individuals are pooled into healthy vs. infected groups, but such approaches typically still use individual-based sequencing to call genotypes before pooling (Wu et al., 2010). When multiple replicates are available, it is possible to combine pool-seq and the case-control design: Endler et al. (2016) analyzed the genetic differences underlying abdominal pigmentation variation among *Drosophila* populations, while Stocks et al. (2019) identified SNPs associated with low versus high ash dieback damage in *Fraxinus excelsior* L. using this approach. Recently, Singh et al. (2024) used a case-control pool-seq approach similar to that deployed here to search for signatures of resistance to Swiss needle cast in *Pseudotsuga menziesii* (Mirb.) Franco. Pool-seq is a cost-effective alternative approach to sequencing of individuals, but the weakness and limitations are also evident, such as unequal representation of individuals and suboptimal allele frequency estimate when the pool size is small, as well as misaligned mapping and sequencing errors (Schlötterer et al., 2014). Nonetheless, in a study on conifer genotyping, Lind et al. (2022) found that allele frequencies estimated from pooled DNA sequencing samples were highly correlated with frequencies estimated from individual sequencing.

Another factor in understanding the genetics of resistance is whether there is interplay between any alleles conferring resistance and introgression between the species. Bechsgaard et al. (2017) showed that plant R genes can adaptively introgress between closely related species. The LP × JP hybrid zone has been very well studied by Cullingham et al. (2012), who found that this hybrid zone presents a mosaic zone with variable introgression and patchy distributions of hybrids (Burns et al., 2019). While it is clear that these species share a broad hybrid zone, it is unclear whether any alleles for disease resistance will be more or less introgressed than the average region in the genome.

In this study, our aim is to identify the genetic basis of DNB resistance in LP, JP, and LP × JP samples and to explore spatial patterning in any identified loci. The specific objectives of our study were to: 1) develop a GWAS case-control approach using pool-seq samples to identify loci with consistent associations with *D. septosporum* response across replicates; and 2) compare the genetic architecture underlying DNB resistance within the studied pine trees and infer how introgression may affect the genetic basis of *D. septosporum* responses. This study provides an important step towards identifying candidate genes to develop genomic tools for screening trees resistant to DNB infection.

## 2 METHODS

### 2.1 Plant materials

Seeds were obtained from 40 natural seedlots across Western Canada (Figure 1, seedlot numbers and locations can be found in Table S1, seed contributors http://adaptree.forestry.ubc.ca/seed-contributors/), including 25 LP seedlots from British Columbia (BC_LP) and three from Alberta (AB_LP), seven LP × JP seedlots from Alberta (AB_LPxJP), and five JP seedlots from Alberta (AB_JP). The range maps were downloaded from https://sites.ualberta.ca/~ahamann/data/rangemaps.html (Hamann et al., 2005). When the seeds were collected in the wild, they were assigned to pure LP, pure JP, or LP × JP, based on their location and morphological traits such as cone and branch characteristics, microfibril angle, and cell area (Wheeler & Guries, 1987; Wood et al., 2009; Yeatman & Teich, 1969). The proportion of LP and JP ancestry of the collected seeds were genotyped using 11 microsatellite loci by Cullingham et al. (2012). Briefly, seeds were germinated to obtain seedlings, then DNA was isolated from the seedlings. DNA was used to amplify 11 microsatellite loci and allele sizes were determined for genotyping as described by Cullingham et al. (2011).

**FIGURE 1.**
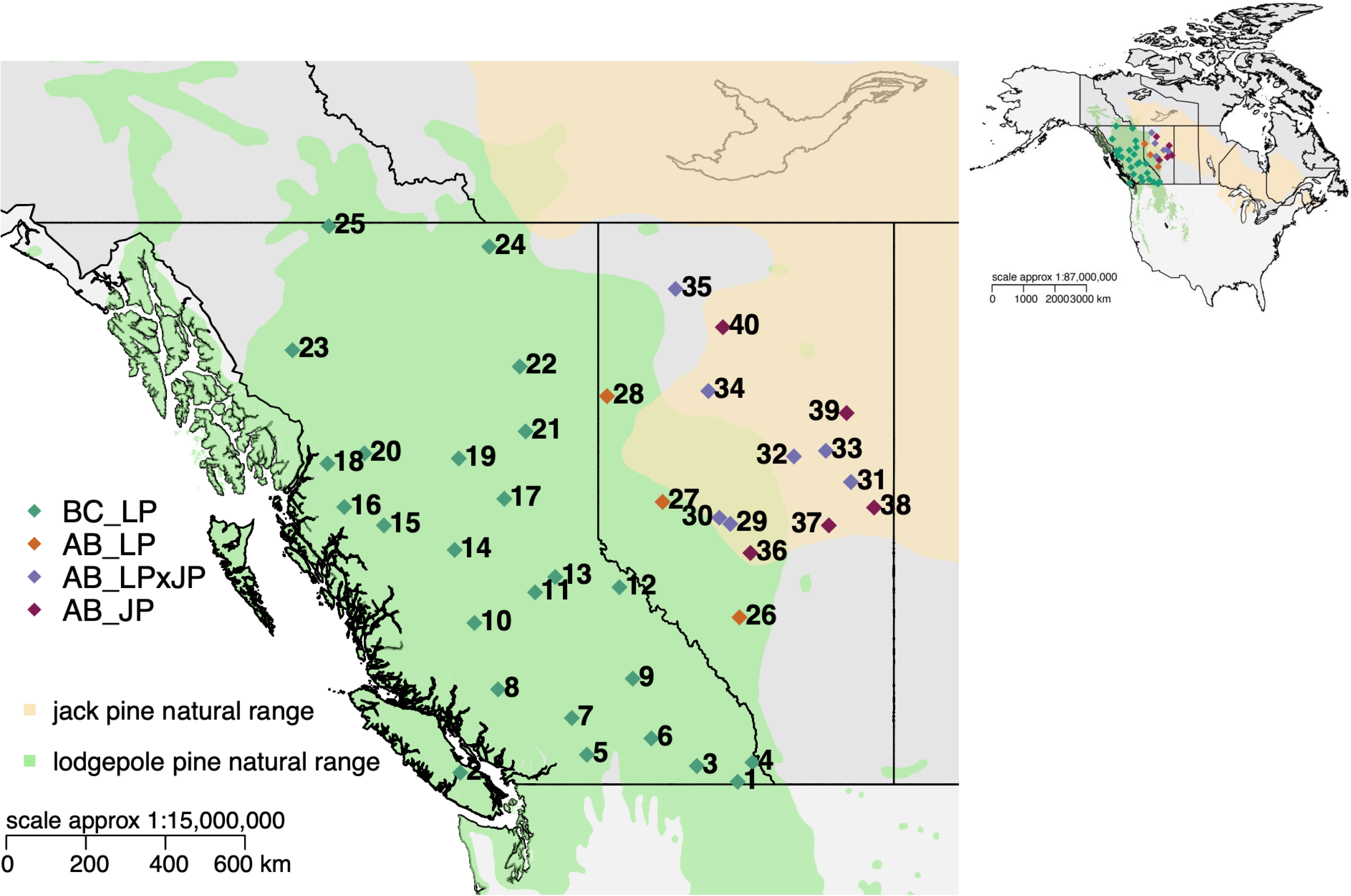
Sampling locations (Table S1) of the 40 pine seedlots analyzed in this study, plotted within the natural range of *Pinus contorta* (lodgepole pine) and *Pinus banksiana* (jack pine) in Canada and USA. The topright map shows the context of the sampling locations. The range maps were downloaded from https://sites.ualberta.ca/~ahamann/data/rangemaps.html. BC_LP (green diamonds) − *Pinus contorta* seedlots from British Columbia (N=25); AB_LP (orange diamonds) − *Pinus contorta* seedlots from Alberta (N=3); AB_ LPxJP (purple diamonds) − *Pinus contorta* × *P. banksiana* hybrids seedlots from Alberta (N=7); AB_JP (brown diamonds) − *Pinus banksiana* seedlots from Alberta (N=5).

For the present study, seeds from the 40 seedlots were used to grow seedlings in a greenhouse at University of British Columbia, Vancouver, BC, for their first year (a flow chart of experimental procedure is shown in Figure S1). Resistance was phenotyped by inoculating 100 individuals per seedlot with one of two *D. septosporum* isolates (D1 & D2, 50 individuals of each isolate). D1 was isolated from needles of an infected LP × JP seedlot in Alberta, Canada, while D2 was isolated from an infected LP seedlot in the Kispiox Valley region close to Smithers in Northwestern British Columbia, Canada (Feau et al., 2021; Ramsfield et al., 2021). The seedlots were identified as LP or LP × JP as aforementioned (Cullingham et al., 2012). The controlled inoculation experiment was performed as described in Kabir et al. (2013) and Feau et al. (2021). Briefly, the one-year-old seedlings were placed in a completely randomized experimental design in a growth chamber with the condition of 16 hours daylight at 20 °C, 8 hours a night at 12 °C, and minimum relative humidity of 80 %. *D. septosporum* conidia were harvested from colonies grown for 10∼16 days on *Dothistroma* sporulation medium plates and then suspended in sterile distilled water. Each seedling was sprayed twice at 10-day intervals with a standard inoculum of approximately 3 mL of 1.6 × 10^6^ conidia mL^−1^, using a household trigger action atomizer. Once the needles were dry (∼45 minutes), the seedlings were wrapped in transparent plastic bags and kept in the growth chamber. After 48h the plastic bags were removed and the seedlings were kept in the growth chamber for 15 weeks till phenotyping. A mister spraying tap water was activated hourly for 3 minutes during the trial to maintain needle wetness. Control seedlings, at least one for each seedlot, were inoculated with sterile water.

Fifteen weeks after inoculation, seedlings were rated for proportion of necrotic needles with red bands and/or fruiting bodies according to a disease severity scale of 1 to 5 (1 = least, 5 = most disease severity). Following disease rating, the ten most- and ten least-infected individuals per seedlot (five individuals for each *D. septosporum* isolate) were identified and retained for DNA extraction. Genomic DNA of each of these individuals was extracted as described in Lind et al. (2022), using the Nucleospin 96 Plant II Core kit (Macherey–Nagel GmbH & Co. KG, Germany) on an Eppendorf epMotion 5075 liquid-handling platform. The DNA of five most-infected and five least-infected individuals by each of the two *D. septosporum* isolates was combined in equimolar amounts to compose four pooled libraries per population, two susceptible (D1-S, D2-S) and two resistant libraries (D1-R, D2-R). Hence,160 pooled DNA libraries were generated.

### 2.2 Sequence capture and pool-seq genotyping

The capture probes were comprised of two sets of probes. The first set of probes was designed based on an existing LP sequence capture array (Suren et al., 2016) by removing the probes that did not yield successful genotyping in Yeaman et al. (2016). Probe sequences of the existing capture array were aligned to the reference genome using GMAP v2019-03-15 (Wu & Watanabe, 2005). Probes that covered the genomic regions with called SNPs in the dataset from Yeaman et al. (2016) were retained, otherwise the probes were discarded. Since there is no available LP reference genome, a masked *Pinus taeda* reference genome Pita.2_01.masked3k2.fa (https://treegenesdb.org/FTP/Genomes/Pita/v2.01/genome/) (Neale et al., 2014) was used instead. *Pinus taeda* is a closely related species to LP (Jin et al., 2021). The second set of probes was newly designed probes derived from the *D. septosporum*-induced genes, which were based on a LP reference transcriptome assembled using the RNA-seq data of *D. septosporum* infected LP samples (Lu et al., 2021). To avoid duplicates, only those *D. septosporum-*induced genes with low homology to the retained probe sequences were used to design the new probes. To do so, the retained probe sequences were aligned to this transcriptome using blastn v2.9.0 with an E-value of 1e-10. A total of 8,778 *D. septosporum*-induced genes did not have any aligned probe sequences. These non-duplicate *D. septosporum-*induced genes were subsequently aligned to the reference genome to predict the exon-intron boundaries using GMAP v2019-03-15. Exon sequences from these induced genes with a length of at least 100 bp were combined with the previously designed working probe sequences, and this combined set of sequences was submitted to Roche NimbleGen (Roche Sequencing Solutions, Inc., CA USA) for Custom SeqCap EZ probe design (design name: 180321_lodgepole_v2_EZ). Combining the two sets of probes, this updated LP sequence capture array has a capture space of 44 Mbp, containing roughly 35,467 assembled genes. Most LP genes responsive to environment stress and fungal pathogen attack were included in the current capture probe design. Though genes expressed in different development periods might be missed, genes that have evidence of substantial expression have been covered in this capture probe design.

The capture libraries for each of the 160 pools were constructed following NimbleGen SeqCap EZ Library SR User’s Guide and as described in Lind et al. (2022). Then the R (resistant) and S (susceptible) libraries (two libraries per capture, R1+S1 or R2+S2, indexed with different barcodes) per population and per isolate were combined for sequence capture and enrichment. Sequencing was performed using the Illumina NovaSeq 6000 S4 PE 150 platform in Centre d’expertise et de services Génome Québec. Our in-house pool-seq pipeline (Lind, 2021) was employed to align the reads to the reference genome and call SNPs. For raw SNPs, only bi-allelic loci in regions without annotated repetitive elements or potentially paralogous genes were retained. The annotated repetitive elements were acquired from the LP genome annotation (Wegrzyn et al., 2014). The potentially paralogous genes were identified as described in Lind et al. (2022) using haploid megagametophyte sequences, for the heterozygous SNP calls for haploid sequences are likely to represent misalignments of paralogs. Afterwards, the SNP loci with depth (DP) < 10, DP > 400, global minor allele frequency < 0.05, or > 25% missing data were also removed.

### 2.3 Genetic structure analyses

The genetic structure among the 160 pooled samples, which represented four pooled libraries (D1-R, D1-S, D2-R, D2-S) for each of the 40 seedlots, was detected by using principal component analysis (PCA), reconstructing a phylogenetic tree, and by assessing correlation of allele frequencies and population differentiation (*F*_ST_) between samples using unlinked SNPs. To reduce the impact of linkage disequilibrium on estimation, the SNP set was thinned using vcftools (Danecek et al., 2011), so that no two sites were within 100,000 bp. PCA and genetic distance were calculated using the major allele depth and the R package “adegenet” (Jombart, 2008). An unrooted phylogenetic tree was constructed using the distance output and the Neighbour-Joining algorithm, which was implemented by the R package “ape” (Paradis & Schliep, 2019). A bootstrap analysis was performed using 5000 bootstrap replicates. Correlation coefficients of major allele frequencies between all pairs within the 160 samples were calculated and plotted using the R package “corrplot” (Wei & Simko, 2017). *F*_ST_ was estimated between all pairs within the 160 samples using the R package “poolfstat” (Hivert et al., 2018). The clustering patterns output from these genetic structure analyses can be used to judge the subtle population structure leading to false positive problems in GWAS.

### 2.4 Dissection of genetic architecture underlying *D. septosporum* response

The alleles and candidate genes associated with *D. septosporum* response were detected using the pool-GWAS method. A GWAS case-control approach was developed using pool-seq samples to identify allele frequency differences between susceptible and resistant pines inoculated with *D. septosporum*. This method was based on Cochran-Mantel-Haenszel (CMH) method and implemented in our in-house pipeline (Lind, 2021). While sequencing read depth was commonly used as an estimation of allele count in such applications of test (Futschik & Schlötterer, 2010; Schlötterer et al., 2014), reliable results would be only achieved when the haploid size of the sample is much greater than the depth of coverage (so that most reads arise from uniquely sampled haplotypes). As the pool size used in the present study (diploid size of 2N=10) is often smaller than the sequencing depth of coverage, read counts were converted into allele frequencies and then multiplied by the ploidy to yield an approximate count for each pool that represents the real replication level. A simulation was conducted to compare the false positive rate from the CMH test using uncorrected or corrected allele counts across a range of number of demes, individuals, and depths. Results showed that the false positive rate of the CMH method is high, especially when the sequencing depth is much higher than the pool size (Figure S2a). Using corrected allele counts, the allele frequencies multiplied by the ploidy (N=10) tends to lower the false positive rate (Figure S2b). Thus, the corrected allele counts were used for CMH test.

As the genetic structure analysis showed that LP × JP and JP samples tend to cluster, JP samples and LP × JP samples were combined (JP + LP × JP) for downstream analyses. Pool-GWAS was performed separately within LP and JP + LP × JP samples. Analysis of R and S samples was conducted by combining individuals inoculated with D1 or D2 isolates to maximize statistical power. To identify candidate genes, SNPs in the top 1% of −log_10_(*p*) values from a pool-GWAS test were classified as outliers. As linkage disequilibrium can amplify signatures of selection, candidate resistance genes were identified as those with a large number of outlier SNPs relative to the average genome-wide expectation, as represented by an index based on the binomial distribution as per Yeaman et al. (2016). The identified loci on the enriched genes were realigned to the 12 LP linkage groups (MacLachlan et al., 2021) and Manhattan plots were drawn. The LP transcriptome (Lu et al., 2021) was used to annotate the reference genome. The aligned transcripts to the reference genome were identified as those with a minimum alignment identity of 90 % and a minimum alignment coverage of 85 % using GMAP v2019-03-15.

### 2.5 Linkage disequilibrium (LD) between loci

A non-linear model (Hill & Weir, 1988) was used to estimate the decay of LD with physical distance. Pairwise correlation (*r*^*2*^) of allele frequencies between loci on the same scaffold were calculated. The *r*^*2*^ values and the physical distances between loci on all scaffolds (genome-wide) or on scaffolds containing the identified top candidate genes (significant) were used to fit the non-linear model as described by Marroni et al. (2011). The LD decay graph was plotted using R (R Core Team, 2018).

### 2.6 Introgression

To study patterns of introgression affecting different regions of the genome and among different samples, the mean pairwise *F*_ST_ was calculated for each locus using the R package “poolfstat”. Loci with high *F*_ST_ values have high genetic differentiation, which may represent species barriers or genomic regions under divergent selection, while loci with low *F*_ST_ values imply introgression. To estimate genetic similarity of the studied JP and LP seedlots, the five JP samples (p36 to p40) and six pure LP samples (the six westmost LP samples, p15, p16, p18, p20, p23, p25) were used to calculate *F*_ST_ values, which were then averaged. To evaluate whether regions associated with *D. septosporum* response have atypical patterns of introgression, the *F*_ST_ values of the top 1 % *D. septosporum* response outliers identified from LP (N = 3,358) and those from JP + LP × JP (N = 3,041) were compared, with *F*_ST_ values of randomly drawn SNPs (N = 3,400).

Similarly, *F*_ST_ between the JP + LP × JP samples (p29 to p40) and six pure LP samples (p15, p16, p18, p20, p23, p25) were averaged to estimate genetic similarity of the studied JP + LP × JP and LP. Different genomic regions were represented by different SNP sets, including *D. septosporum* response outliers identified from LP (N = 3,358) and outliers identified from JP + LP × JP (N = 3,041), as well as the unlinked SNP set (N = 31,716), which was used in 2.3 for detecting genetic structure and representing random genomic region. The genetic similarity (*F*_ST_) of JP + LP × JP and LP was regressed to the longitude of these JP + LP × JP samples. The regression slopes for the three SNPs sets were compared using analysis of covariance method implemented by the R package “lsmeans” (Lenth, 2016; R Core Team, 2018).

## 3 RESULTS

### 3.1 Patterns of genetic structure

After filtering, a total of 364,691 SNP loci were retained for downstream analyses, and most loci had a minor allele frequency between 0.05 and 0.1 (Figure S3). A thinned set of 31,716 unlinked SNPs was used for studying genetic structure among the 160 pooled samples (Figure 2). The most prominent patterns of structure were associated with species (Figure 2a & b, Figure S4), with LP samples readily distinguishable from JP and LP × JP samples. LP × JP samples clustered more closely with JP than LP. Compared to those sampled from the eastern hybrid zone, western LP × JP tend to be more similar to LP (samples were arranged according to longitude from left to right in Figure 2 c & d). Given this clear pattern of genetic structure, we conducted GWAS in LP and JP + LP × JP samples separately.

**FIGURE 2.**
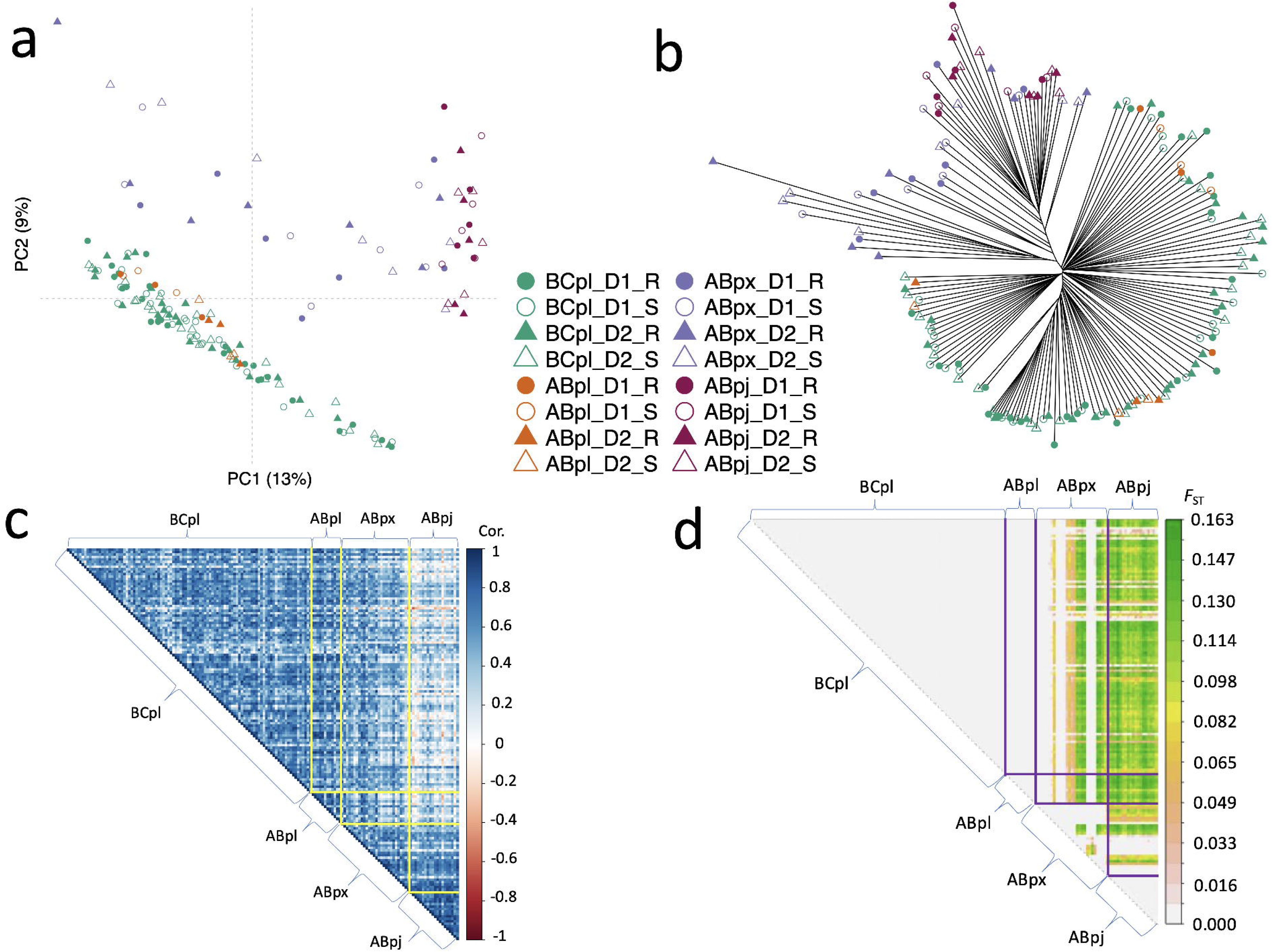
Genetic structure among the 160 pooled samples. a) Principal component analysis. b) Unrooted phylogenetic tree. The bootstrap values for each branch were shown in Figure S4. c) Correlation plot of the major allele frequencies between samples. d) Pairwise *F*_ST_ values between pools. BC_LP − *Pinus contorta* (lodgepole pine) samples from British Columbia; AB_LP − *Pinus contorta* (lodgepole pine) samples from Alberta; AB_LPxJP − *Pinus contorta* × *P. banksiana* hybrids samples from Alberta; AB_JP − *Pinus banksiana* (jack pine) samples from Alberta. D1 − *D. septosporum* isolate 1; D2 − *D. septosporum* isolate 2. R − resistant tree; S − susceptible tree. In c) and d), each column represents a pooled sample. The samples were arranged from left to right according to the longitude of sampling location, and four pooled libraries per seedlot (D1-R, D1-S, D2-R, D2-S) were arranged together.

### 3.2 Signatures of association to infection response

Out of the 100 top-ranked candidate genes in each test (Table S2), three were identified within both the LP and JP + LP × JP samples (Figure 3). This is significantly more than the hypergeometric expectation, based on the probability of overlap for two draws of 100 genes from 20,026 genes. A previous study anchored 10,093 scaffolds on the LP linkage map (MacLachlan et al., 2021), so we plotted SNPs and labeled the enriched plant-pathogen interaction candidate genes on the 12 linkage groups (Figure 3). The three genes identified in both species encode ABC transporter, F-box Kelch-repeat protein, and serine/threonine protein kinase. Genes encoding F-box protein and serine protein kinase were highlighted in Figure 3. Genes encoding ABC transporter were not included in the linkage groups, so they were not included in Figure 3. We also checked the enrichment of differentially expressed genes (DEGs) in the top 100 ranked genes. Out of 631 DEGs identified in the previous study (Lu et al., 2021), four were also candidate genes in the present study. Fisher’s Exact test showed there is no significant enrichment of DEGs in the candidate genes.

**FIGURE 3.**
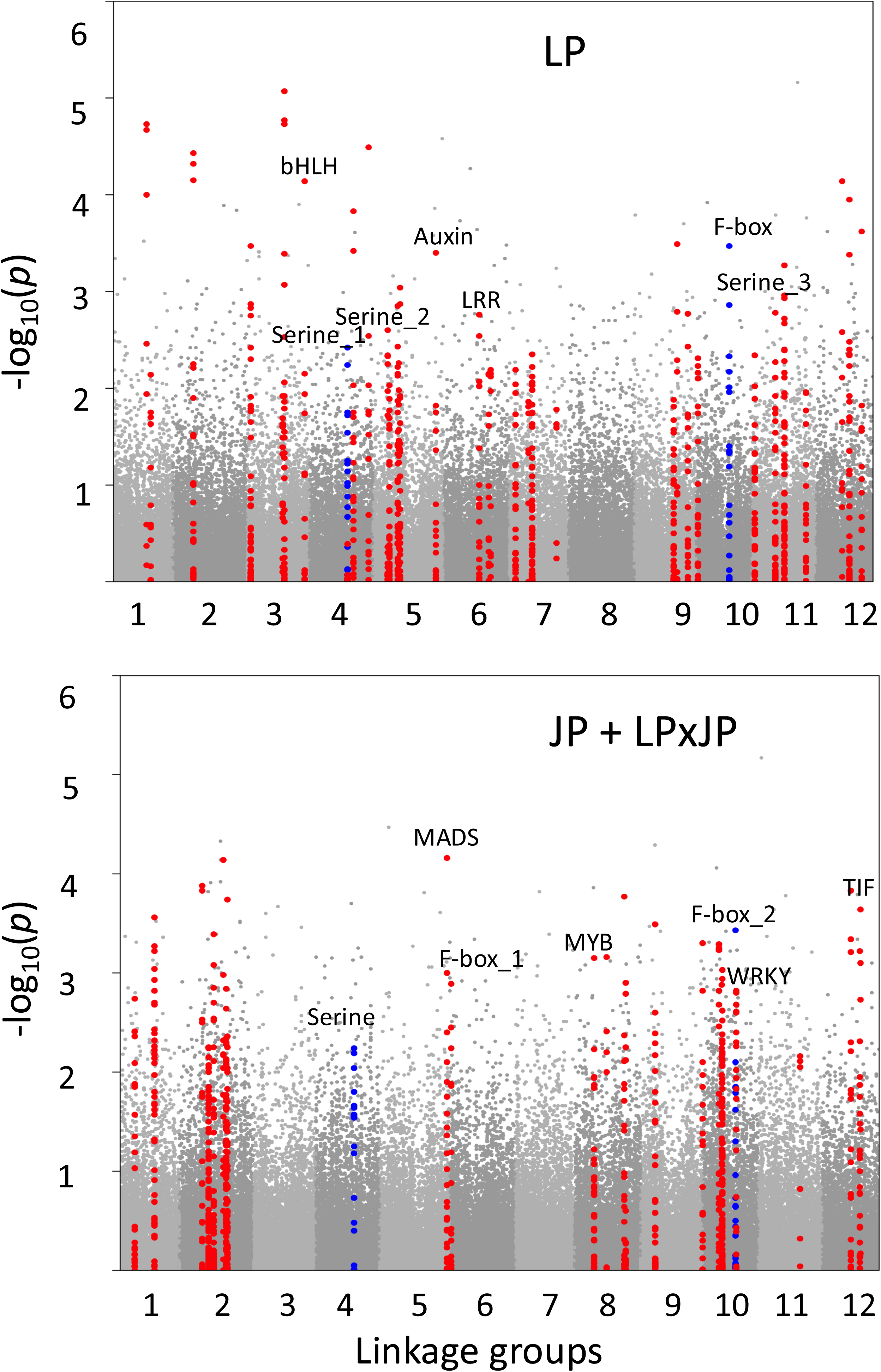
Manhattan plot showing SNPs associated with *D. septosporum* response within *Pinus contorta* (lodgepole pine, LP) and combined *Pinus banksiana* and *Pinus contorta* × *P. banksiana* hybrids (JP + LPxJP) samples. The x axis represents 12 linkage groups. The y axis represents significance of associations identified from pool-GWAS. Candidate genes were identified using a binomial test for enrichment of outlier SNPs per gene. Candidate genes and their surrounding 50 kbp regions are highlighted in red. Of the three top-ranked candidate genes identified within both the LP and JP + LPxJP samples, one was not present in the linkage map, while the other two are highlighted in blue (“Serine” on linkage group 4 and “F-box” on linkage group 10). Labels in the graphs represent candidate genes encoding plant-pathogen interaction proteins: bHLH − bHLH transcription factor; Serine − serine/threonine-protein kinase; Auxin − auxin response factor; LRR − proteins with leucine-rich repeat; F-box − F-box Kelch-repeat protein; MADS − MADS transcription factor; MYB − MYB transcription factor; WRKY − WRKY transcription factor; TIF − translation initiation factor. The candidate gene IDs (in Table S2) are: In LP, bHLH − Scaffold_2015-465162-569336; Serine_1 − Scaffold_3728-494595-736720; Serine_2 − Scaffold_3141-1-105566; Serine_3 − Scaffold_1072-1299306-1374553; Auxin − Scaffold_3823-1571412-1689102; LRR − Scaffold_1361-487806-570406; F-box − Scaffold_3698-733755-839782; In JP + LPxJP, Serine − Scaffold_3728-494595-736720; MADS − Scaffold_427-2324701-2717580; F-box_1 − Scaffold_3698-733755-839782; MYB − Scaffold_4478-441537-627389; F-box_2 − Scaffold_342-1965787-2253515; WRKY − Scaffold_830-1136302-1239031; TIF − Scaffold_207-7583953-7695067.

Loci on scaffolds with outliers (the SNPs with top 1 % of −log_10_(*p*) values from pool-GWAS test) tend to exhibit a more gradual decay in LD compared with the genome-wide loci on all scaffolds (Figure 4), with JP + LP × JP pine outliers having a much slower decay rate than LP. In LP, the distance at which LD decays to half of its maximum value is 96 bp for genome-wide loci, and 174 bp for outlier loci. In JP + LP × JP, the half LD decay distance is 1,357 bp for genome-wide loci, and 68,215 bp for outlier loci. The slower LD decay pattern for the outliers may indicate selection on these identified loci.

**FIGURE 4.**
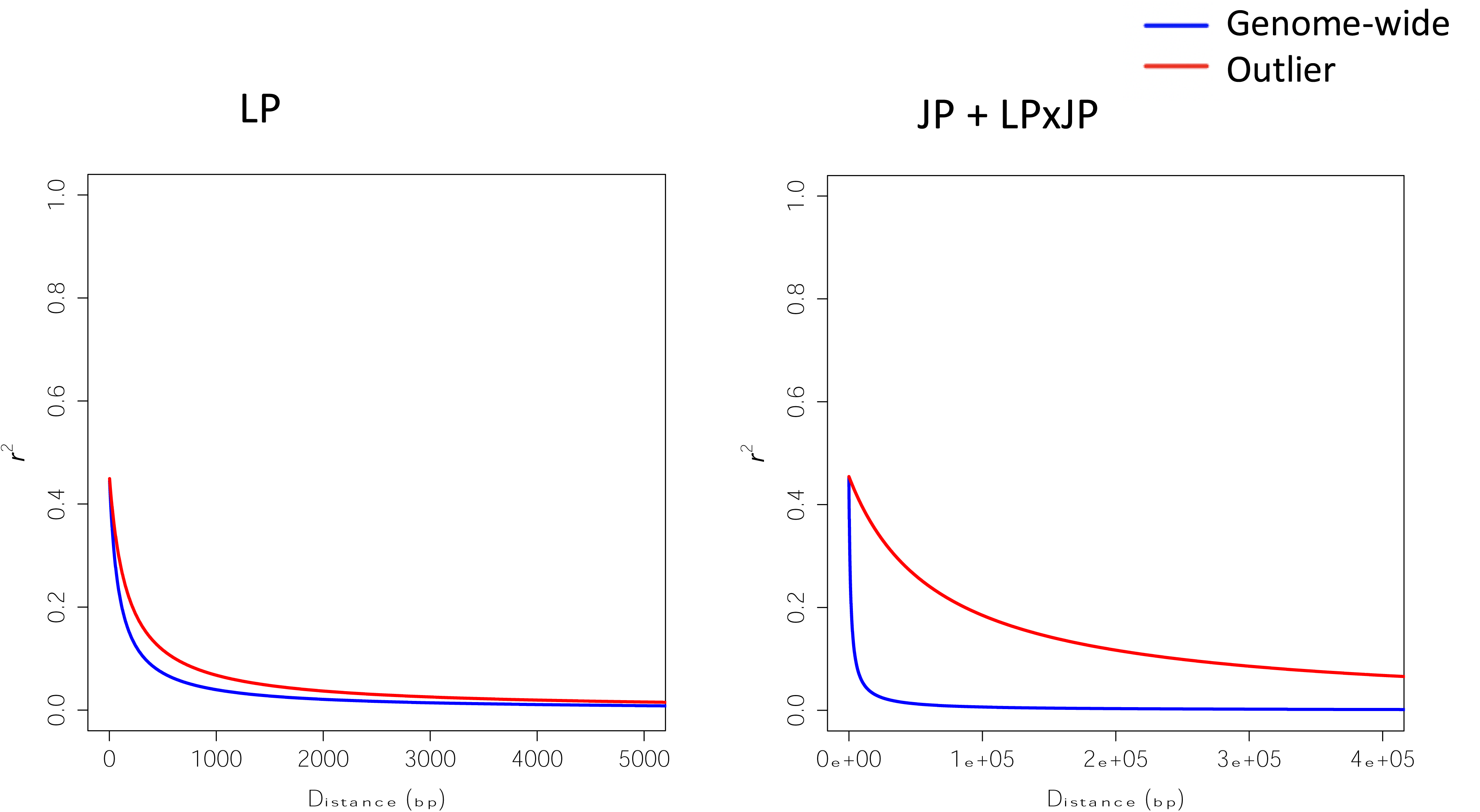
Linkage disequilibrium (LD) decay for genome-wide loci and outlier loci identified from *Pinus contorta* (lodgepole pine, LP) and combined *Pinus banksiana* and *Pinus contorta* × *P. banksiana* hybrids (JP + LPxJP) samples. The *r*^*2*^ values (pairwise correlation of allele frequencies between loci) and the physical distances between loci were used to fit the non-linear model as described by Marroni et al. (2011).

### 3.3 Introgression

By calculating average *F*_ST_ between pairs of LP and JP samples,we found that the *D. septosporum* response outliers, which were identified separately in LP or JP + LP × JP samples, have higher *F*_ST_ values than randomly drawn SNPs (Wilcoxon test *p*-value < 0.01, Figure 5a & 5b), with outliers identified in JP + LP × JP samples showing higher *F*_ST_ values than those identified in LP samples (Wilcoxon test *p*-value < 0.01). These results suggest that the outliers associated with disease response constitute or are linked to genomic regions for divergent selection or species barriers, rather than facilitating introgression. *F*_ST_ estimates between JP + LP × JP samples and a set of pure LP samples showed a pattern of increased differentiation with distance eastwards of the JP + LP × JP samples (Figure 5c), which were higher for the outliers than randomly chosen SNPs. Given the small number of samples, significance of linear model fitting is borderline, with *p*-value = 0.06 for outliers identified in LP, *p*-value = 0.05 for outliers identified in JP + LP × JP, and *p*-value = 0.06 for randomly chosen SNPs (df = 10). The differences between regression slopes on different gene sets (LP outliers, JP + LP × JP outliers, and unlinked SNPs) were not significant (*p*-values of pairwise comparisons > 0.3). Though it is not possible to test whether a linear vs. discontinuous model would fit better to these data, the results suggest that isolation-by-distance is occurring here.

**FIGURE 5.**
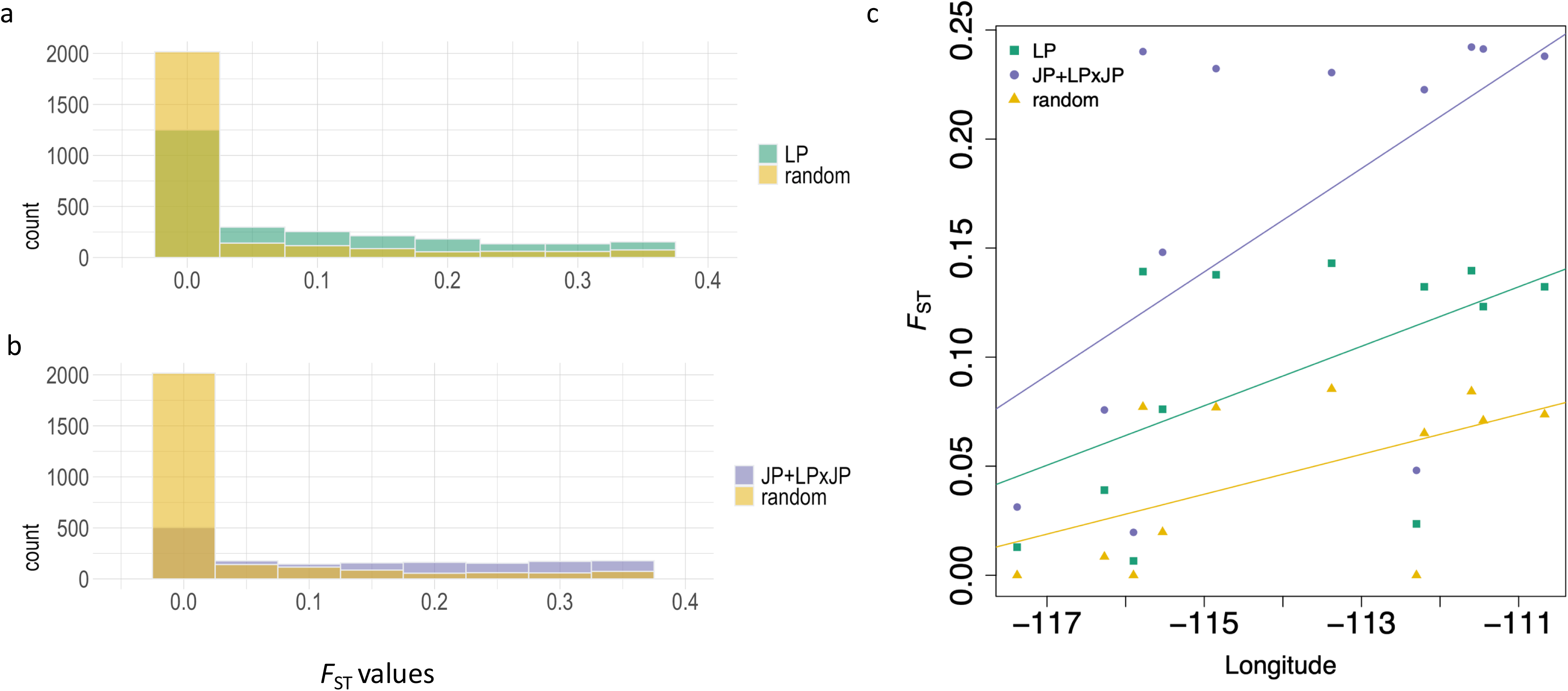
*F*_ST_ values of *D. septosporum* response outliers (a & b) and regression patterns of *F*_ST_ values with longitude. For a & b, the *F*_ST_ values were calculated between the *Pinus banksiana* (jack pine, JP) samples (p36 to p40) and the six pure *Pinus contorta* (lodgepole pine, LP) samples (p15, p16, p18, p20, p23, p25).JP + LPxJP represents the combined JP and *Pinus contorta* × *P. banksiana* hybrids samples. *F*_ST_ values of *D. septosporum* response outliers identified from LP (N=3,358) and JP + LPxJP (N=3,041) samples were compared, with *F*_ST_ values of randomly drawn SNPs (N=3,400). Outliers identified from both species have higher *F*_ST_ values than randomly drawn SNPs (*p*-value < 0.01). For c, the *F*_ST_ values were calculated between the JP + LPxJP samples (p29 to p40) and the six pure LP samples (p15, p16, p18, p20, p23, p25) and then averaged for each JP + LPxJP sample. Different genomic regions were represented by different SNP sets, including LP outliers (N=3,358), JP + LPxJP outliers (N=3,041), and unlinked SNPs (N=31,716). The *F*_ST_ values represented by different SNP sets were regressed to the longitude of these JP + LPxJP samples. Differences between slopes are not significant (ANCOVA, *p*-value > 0.3).

## 4 DISCUSSION

In recent decades, the fungal pathogen *D. septosporum* has become an increasing threat to LP in British Columbia (Feau et al., 2021; Welsh et al., 2009). Although *Dothistroma* needle blight (DNB) has not been observed in natural stands of JP in Alberta, Ramsfield et al. (2021) confirmed JP’s susceptibility to infection by *D. septosporum* using a trial under natural conditions. The potential of climate change to allow this pathogen to spread across the wide natural distribution of JP raises serious concerns. The hybrid zone of contact between the sister species LP and JP may act as a bridge, allowing this fungal pathogen host shift (Taylor et al., 2015), as evidenced by the intermediate phenotype of gall development in LP × JP seedlings compared to that in LP and JP (McAllister et al., 2022). The genetic basis of pathogen resistance in each species is therefore an important factor affecting the potential for host shift. To study the genetics of resistance to DNB, we analyzed the genetic architecture underlying *D. septosporum* response using the studied samples of LP, JP, and LP × JP. We identified candidate genes associated with *D. septosporum* response and inferred the impact of introgression on the genetic structure underlying *D. septosporum* response in the studied samples. We detected largely non-overlapping sets of candidate genes in LP and JP + LP × JP in response to *D. septosporum*, which might be shaped by different evolutionary forces of selection and introgression.

Alternatively, the non-overlapping sets of candidate genes might reflect that many non-causal genes exist in the top 1 % GWAS outliers of each species, given that we did not test for significance at the gene-level. In either case, it seems clear that resistance to DNB is a highly polygenic trait with alleles of small effect.

This study is the first report of genome-wide scanning of *D. septosporum* response genes in LP, JP, and LP × JP. We identified a few top candidate genes encoding leucine-rich repeats (LRR)-containing protein domain and serine threonine kinases in LP samples (Figure 3, Table S2). Plant breeders have long used nucleotide-binding LRR (NB-LRR) receptor R genes for preventing or reducing diseases in crops. When attacked by infectious pathogens, plant NB-LRR receptors can recognize pathogen effectors by either direct or indirect mechanisms (Dodds & Rathjen, 2010). Such strategy is called effector-triggered immunity (ETI), which occurs after a basal resistance, pathogen-associated-molecular-patterns-triggered immunity (PTI). In a previous study using RNA-seq data, Lu et al. (2021) identified genes involved in PTI and ETI, as well as R genes with positive selection signals in LP. These results indicate that the long coevolutionary history between LP and *D. septosporum* (Capron et al., 2021; Welsh et al., 2009) has given rise to some tolerance or resistance mechanism against DNB. Since most of the identified associated markers are likely not causal but rather are in LD with the causal loci, the observed fast LD decay rate in lodgepole pine implies promising fine mapping resolutions as the identified markers are likely to be proximal to the causative genes.

On the contrary, JP, as a potential new host, may not have coevolved such a specific defense, and the long LD blocks surrounding candidate genes identified from JP + LP × JP samples, which are typical of hybrid zones, limit the resolution of association mapping, so it may be difficult to locate the causative genes for follow-up studies (Goulet et al., 2017; Wilson & Goldstein, 2000). Additionally, we caution that the capture probes were designed using LP not JP genes, ascertainment bias might give rise to the underrepresentation of JP alleles (Lachance & Tishkoff, 2013). However, we can use these JP + LP × JP candidate genes to infer how introgression affects the frequencies of disease-resistant alleles. Since we analyzed pool-seq data and lacked the pure JP samples out of the hybrid zone, it is not possible to analyze parental origin information, thus we inferred the ancestry using clustering. The clustering patterns in Figure 2 show that most hybrid pine trees in our study had greater JP than LP ancestry, likely in part due to our limited sampling of JP from further Eastwards. Western LP × JP tend to be more genetically similar to LP, while eastern LP × JP tend to be more genetically dissimilar to LP. This is a typical characteristic for ongoing introgression and hybridization in the contact zone with a possible trend of LP expanding its range eastward (Harrison & Larson, 2014; Moran et al., 2021).

In hybrid zones, alleles in some genomic regions are able to introgress across species boundaries, whereas alleles that constitute species barriers or are under divergent selection will remain differentiated (Harrison & Larson, 2014). We found significantly high *F*_ST_ values between JP and LP samples for the candidate genes associated with *D. septosporum* infection response. This can occur if these candidate genes tend to reside in genomic regions with restricted gene flow, driven by linkage with genes for divergent selection or species barriers. McAllister et al. (2022) observed a gradient of resistance to *C. harknessii* in LP, which could be associated with introgression of resistance genes from JP into LP. Since LP might have a long coevolutionary history with *D. septosporum* (Capron et al., 2021; Welsh et al., 2009), the introgression of LP genes into hybrids may increase resistance to *D. septosporum*. However, variation in *F*_ST_ along longitude (Figure 5c) did not differ significantly between background and candidate loci, so we cannot draw strong conclusions about adaptive introgression here.

## 5 CONCLUSION

The recent outbreaks of DNB in LP and its ability to infect JP pose a pressing need to understanding the genetic architecture of resistance to *D. septosporum*. The present study provides an array of candidate genes associated with *D. septosporum* response using pool-

GWAS. The identified candidate genes within LP and JP + LP × JP samples are largely non-overlapping, and the adaptive introgression is yet to be found. This knowledge can be used to develop genomic tools to minimize DNB risk and guide forest management strategies. Further testing and validation of these associations using a SNP array is currently underway.

## FIGURE LEGENDS

## AUTHOR CONTRIBUTIONS

Mengmeng Lu and Sam Yeaman conceived analyses and wrote the manuscript. Mengmeng Lu performed computational analyses. Nicolas Feau and Richard C. Hamelin designed pathogen inoculation and phenotyping protocol. Nicolas Feau performed the pathogen inoculation and phenotyping in the greenhouse. Brandon Lind and Pooja Singh designed pool-GWAS analyses pipeline. Dragana Obreht Vidakov performed sequence capture work. Sally N. Aitken, Richard C. Hamelin, and Sam Yeaman obtained funding. All authors edited the manuscript.

## ACKNOWLEDGMENTS

We thank Centre d’expertise et de services Génome Québec for sequencing service, University of Calgary Information Technologies for system support, the Research Oversight Committee of CoAdapTree Project for suggestions and help. Special gratitude goes to Dr. Andy Benowicz and our seed contributors http://adaptree.forestry.ubc.ca/seed-contributors/.

## DATA AVAILABILITY STATEMENT

The raw pool-seq data were deposited in NCBI SRA (accession number: PRJNA602898; http://www.ncbi.nlm.nih.gov/sra). The assembled transcriptomes and their annotation were deposited in Dryad https://doi.org/10.5061/dryad.prr4xgxtk. The scripts used for analyses were deposited in https://github.com/Mengmeng-Lu/Genetic-architecture-underlying-response-to-the-fungal-pathogen-Dothistroma-septosporum-in-Pinus-con

## REFERENCES

Alonso-Blanco, C. & Méndez-Vigo, B. (2014). Genetic architecture of naturally occurring quantitative traits in plants: an updated synthesis. Current opinion in plant biology, 18, 37–43. 10.1016/j.pbi.2014.01.002

Bechsgaard, J., Jorgensen, T. H., & Schierup, M. H. (2017). Evidence for adaptive introgression of disease resistance genes among closely related Arabidopsis species. G3 (Bethesda), 7(8), 2677–2683. 10.1534/g3.117.043984

Boroń, P., Lenart-Boroń, A., Mullett, M., Grad, B., & Nawrot-Chorabik, K. (2021). Population structure of Dothistroma septosporum in Poland: revealing the genetic signature of a recently established pathogen. Plant Pathology, 70(6), 1310–1325. 10.1111/ppa.13383

Burns, I., James, P. M. A., Coltman, D. W., & Cullingham, C. I. (2019). Spatial and genetic structure of the lodgepole × jack pine hybrid zone. Canadian Journal of Forest Research, 49(7), 844–853. 10.1139/cjfr-2018-0428

Capador-Barreto, H. D., Bernhardsson, C., Milesi, P., Vos, I., Lundén, K., Wu, H. X., Elfstrand, M. (2021). Killing two enemies with one stone? Genomics of resistance to two sympatric pathogens in Norway spruce. Molecular Ecology, 30(18), 4433–4447. 10.1111/mec.16058

Capron, A., Feau, N., Heinzelmann, R., Barnes, I., Benowicz, A., Bradshaw, R. E., Hamelin, R. C. (2021). Signatures of post-glacial genetic isolation and human-driven migration in the Dothistroma needle blight pathogen in Western Canada. Phytopathology, 111(1), 116–127. 10.1094/phyto-08-20-0350-fi

Cullingham, C. I., Cooke, J. E., Dang, S., Davis, C. S., Cooke, B. J., & Coltman, D. W. (2011). Mountain pine beetle host-range expansion threatens the boreal forest. Molecular Ecology, 20(10), 2157–2171. 10.1111/j.1365-294X.2011.05086.x

Cullingham, C. I., James, P. M., Cooke, J. E., & Coltman, D. W. (2012). Characterizing the physical and genetic structure of the lodgepole pine× jack pine hybrid zone: mosaic structure and differential introgression. Evolutionary Applications, 5(8), 879–891. 10.1111/j.1752-4571.2012.00266.x

Dale, A. L., Lewis, K. J., & Murray, B. W. (2011). Sexual reproduction and gene flow in the pine pathogen Dothistroma septosporum in British Columbia. Phytopathology, 101(1), 68–76. 10.1094/phyto-04-10-0121

Danecek, P., Auton, A., Abecasis, G., Albers, C. A., Banks, E., DePristo, M. A., 1000 Genomes Project Analysis Group. (2011). The variant call format and VCFtools. Bioinformatics, 27(15), 2156–2158. 10.1093/bioinformatics/btr330

De La Torre, A. R., Puiu, D., Crepeau, M. W., Stevens, K., Salzberg, S. L., Langley, C. H., & Neale, D. B. (2019). Genomic architecture of complex traits in loblolly pine. New Phytologist, 221(4), 1789–1801. 10.1111/nph.15535

Dodds, P. N. & Rathjen, J. P. (2010). Plant immunity: towards an integrated view of plant– pathogen interactions. Nature review genetics, 11(8), 539–548. 10.1038/nrg2812

Endler, L., Betancourt, A. J., Nolte, V., & Schlötterer, C. (2016). Reconciling differences in pool-GWAS between populations: A case study of female abdominal pigmentation in Drosophila melanogaster. Genetics, 202(2), 843–855. 10.1534/genetics.115.183376

Feau, N., Ramsfield, T. D., Myrholm, C. L., Tomm, B., Cerezke, H. F., Benowicz, A., Hamelin, R. C. (2021). DNA-barcoding identification of Dothistroma septosporum on Pinus contorta var. latifolia, P. banksiana and their hybrid in northern Alberta, Canada. Canadian Journal of Plant Pathology, 43(3), 472–479. 10.1080/07060661.2020.1829065

Futschik, A. & Schlötterer, C. (2010). The next generation of molecular markers from massively parallel sequencing of pooled DNA Samples. Genetics, 186(1), 207–218. 10.1534/genetics.110.114397

Gibson, I. A. S. (1972). Dothistroma Blight of Pinus Radiata. Annual Review of Phytopathology, 10(1), 51–72. 10.1146/annurev.py.10.090172.000411

Goulet, B. E., Roda, F., & Hopkins, R. (2017). Hybridization in plants: Old ideas, new techniques. Plant physiology, 173(1), 65–78. 10.1104/pp.16.01340

Hall, D. E., Yuen, M. M. S., Jancsik, S., Quesada, A. L., Dullat, H. K., Li, M., Bohlmann, J. (2013). Transcriptome resources and functional characterization of monoterpene synthases for two host species of the mountain pine beetle, lodgepole pine (Pinus contorta) and jack pine (Pinus banksiana). BMC plant biology, 13(1), 80. 10.1186/1471-2229-13-80

Hamann, A., Smets, P., Yanchuk, A. D., & Aitken, S. N. (2005). An ecogeographic framework for in situ conservation of forest trees in British Columbia. Canadian Journal of Forest Research, 35(11), 2553–2561. 10.1139/x05-181

Hao, Z.-Z., Liu, Y.-Y., Nazaire, M., Wei, X.-X., & Wang, X.-Q. (2015). Molecular phylogenetics and evolutionary history of sect. Quinquefoliae (Pinus): Implications for Northern Hemisphere biogeography. Molecular Phylogenetics and Evolution, 87, 65–79. 10.1016/j.ympev.2015.03.013

Harrison, R. G. & Larson, E. L. (2014). Hybridization, introgression, and the nature of species boundaries. Journal of Heredity, 105(S1), 795–809. 10.1093/jhered/esu033

Hill, W. G. & Weir, B. S. (1988). Variances and covariances of squared linkage disequilibria in finite populations. Theoretical population biology, 33(1), 54–78. 10.1016/0040-5809(88)90004-4

Hivert, V., Leblois, R., Petit, E. J., Gautier, M., & Vitalis, R. (2018). Measuring genetic differentiation from pool-seq data. Genetics, 210(1), 315–330. 10.1534/genetics.118.300900

Isabel, N., Holliday, J. A., & Aitken, S. N. (2019). Forest genomics: Advancing climate adaptation, forest health, productivity, and conservation. Evolutionary Applications, 13(1), 3–10. 10.1111/eva.12902

Jia, H., Zhang, Y., Orbović, V., Xu, J., White, F. F., Jones, J. B., & Wang, N. (2017). Genome editing of the disease susceptibility gene CsLOB1 in citrus confers resistance to citrus canker. Plant biotechnology journal, 15(7), 817–823. 10.1111/pbi.12677

Jin, W.-T., Gernandt, D. S., Wehenkel, C., Xia, X.-M., Wei, X.-X., & Wang, X.-Q. (2021). Phylogenomic and ecological analyses reveal the spatiotemporal evolution of global pines. Proceedings of the National Academy of Sciences, 118(20), e2022302118. 10.1073/pnas.2022302118

Jombart, T. (2008). adegenet: a R package for the multivariate analysis of genetic markers. Bioinformatics, 24(11), 1403–1405. 10.1093/bioinformatics/btn129

Kabir, M. S., Ganley, R. J., & Bradshaw, R. E. (2013). An improved artificial pathogenicity assay for Dothistroma needle blight on Pinus radiata. Australasian Plant Pathology, 42(4), 503–510. 10.1007/s13313-013-0217-z

Lachance, J. & Tishkoff, S. A. (2013). SNP ascertainment bias in population genetic analyses: why it is important, and how to correct it. Bioessays, 35(9), 780–786. 10.1002/bies.201300014

Lenth, R. V. (2016). Least-squares means: The R package lsmeans. Journal of Statistical Software, 69(1), 1–33. 10.18637/jss.v069.i01

Lind, B. M. (2021). GitHub.com/brandonlind/cmh_test: preprint release (Version 1.0.0). Zenodo. 10.5281/zenodo.5083798

Lind, B. M. (2021). GitHub.com/CoAdapTree/varscan_pipeline: Publication release (Version 1.0.0). Zenodo. 10.5281/zenodo.5083302

Lind, B. M., Lu, M., Obreht Vidakovic, D., Singh, P., Booker, T. R., Aitken, S. N., & Yeaman, S. (2022). Haploid, diploid, and pooled exome capture recapitulate features of biology and paralogy in two non-model tree species. Molecular Ecology Resources, 22(1), 225–238. 10.1111/1755-0998.13474

Lu, M., Feau, N., Obreht Vidakovic, D., Ukrainetz, N., Wong, B., Aitken, S. N., Yeaman, S. (2021). Comparative gene expression analysis reveals mechanism of Pinus contorta response to the fungal pathogen Dothistroma septosporum. Molecular plant-microbe interactions: MPMI, 34(4), 397–409. 10.1094/mpmi-10-20-0282-r

Lu, M., Krutovsky, K. V., Nelson, C. D., West, J. B., Reilly, N. A., & Loopstra, C. A. (2017). Association genetics of growth and adaptive traits in loblolly pine (Pinus taeda L.) using whole-exome-discovered polymorphisms. Tree Genetics & Genomes, 13(3), 57. 10.1007/s11295-017-1140-1

MacLachlan, I. R., McDonald, T. K., Lind, B. M., Rieseberg, L. H., Yeaman, S., & Aitken, S. N. (2021). Genome-wide shifts in climate-related variation underpin responses to selective breeding in a widespread conifer. Proceedings of the National Academy of Sciences, 118(10), e2016900118. 10.1073/pnas.2016900118

Marroni, F., Pinosio, S., Zaina, G., Fogolari, F., Felice, N., Cattonaro, F., & Morgante, M. (2011). Nucleotide diversity and linkage disequilibrium in Populus nigra cinnamyl alcohol dehydrogenase (CAD4) gene. Tree Genetics & Genomes, 7(5), 1011–1023. 10.1007/s11295-011-0391-5

McAllister, C. H., Cullingham, C. I., Peery, R. M., Mbenoun, M., McPeak, E., Feau, N., Cooke, J. E. K. (2022). Evidence of coevolution between Cronartium harknessii lineages and their corresponding hosts, lodgepole pine and jack pine. Phytopathology, 112(8), 1795–1807. 10.1094/phyto-09-21-0370-r

Min, T., Hwarari, D., Li, D., Movahedi, A., & Yang, L. (2022). CRISPR-based genome editing and its applications in woody plants. International journal of molecular sciences, 23(17), 10175. 10.3390/ijms231710175

Moran, B. M., Payne, C., Langdon, Q., Powell, D. L., Brandvain, Y., & Schumer, M. (2021). The genomic consequences of hybridization. eLife, 10, e69016. 10.7554/eLife.69016

Neale, D. B., Wegrzyn, J. L., Stevens, K. A., Zimin, A. V., Puiu, D., Crepeau, M. W., Langley, C. H. (2014). Decoding the massive genome of loblolly pine using haploid DNA and novel assembly strategies. Genome Biology, 15(3), R59. 10.1186/gb-2014-15-3-r59

Paradis, E. & Schliep, K. (2019). ape 5.0: an environment for modern phylogenetics and evolutionary analyses in R. Bioinformatics, 35(3), 526–528. 10.1093/bioinformatics/bty633

Peery, R. M., McAllister, C. H., Cullingham, C. I., Mahon, E. L., Arango-Velez, A., & Cooke, J. E. K. (2021). Comparative genomics of the chitinase gene family in lodgepole and jack pines: contrasting responses to biotic threats and landscape level investigation of genetic differentiation. Botany, 99(6), 355–378. 10.1139/cjb-2020-0125

Quesada, T., Gopal, V., Cumbie, W. P., Eckert, A. J., Wegrzyn, J. L., Neale, D. B., Davis, J. M. (2010). Association mapping of quantitative disease resistance in a natural population of loblolly pine (Pinus taeda L.). Genetics, 186(2), 677–686. 10.1534/genetics.110.117549

R Core Team. (2018). R: A language and environment for statistical computing. Vienna, Austria: R Foundation for Statistical Computing. https://www.R-project.org/.

Ramsfield, T., Myrholm, C., & Tomm, B. (2021). Natural infection of Pinus contorta var. latifolia, Pinus banksiana and their hybrid by Dothistroma septosporum in Alberta, Canada. Forest Pathology, 51(5), e12717. 10.1111/efp.12717

Rudolph, T. D. & Laidly, P. R. (1990). Pinus banksiana Lamb. Jack Pine. In R. M. Burns & B. H. Honkala (Eds.), Silvics of North America (Vol. Volume 1: Conifers. Agriculture Handbook 654, pp. 280–293). Washington, D.C.: USDA Forest Service.

Rudolph, T. D. & Yeatman, C. W. (1982). Genetics of jack pine. In Research Paper WO-38 (pp. 35–37). Washington, D.C.: USDA Forest Service.

Schlötterer, C., Tobler, R., Kofler, R., & Nolte, V. (2014). Sequencing pools of individuals — mining genome-wide polymorphism data without big funding. Nature Reviews Genetics, 15(11), 749–763. 10.1038/nrg3803

Seidl, R., Thom, D., Kautz, M., Martin-Benito, D., Peltoniemi, M., Vacchiano, G., Reyer, C. P. O. (2017). Forest disturbances under climate change. Nature Climate Change, 7(6), 395–402. 10.1038/nclimate3303

Simler-Williamson, A. B., Rizzo, D. M., & Cobb, R. C. (2019). Interacting effects of global change on forest pest and pathogen dynamics. Annual Review of Ecology, Evolution, and Systematics, 50, 381–403. 10.1146/annurev-ecolsys-110218-024934

Singh, P., St Clair, J. B., Lind, B. M., Cronn, R., Wilhelmi, N. P., Feau, N., Yeaman, S. (2024). Genetic architecture of disease resistance and tolerance in Douglas-fir trees. New Phytologist, 243(2), 705–719. 10.1111/nph.19797

Stocks, J. J., Metheringham, C. L., Plumb, W. J., Lee, S. J., Kelly, L. J., Nichols, R. A., & Buggs, R. J. A. (2019). Genomic basis of European ash tree resistance to ash dieback fungus. Nature Ecology & Evolution, 3(12), 1686–1696. 10.1038/s41559-019-1036-6

Suren, H., Hodgins, K. A., Yeaman, S., Nurkowski, K. A., Smets, P., Rieseberg, L. H., Holliday, J. A. (2016). Exome capture from the spruce and pine giga-genomes. Molecular Ecology Resources, 16(5), 1136–1146. 10.1111/1755-0998.12570

Tam, V., Patel, N., Turcotte, M., Bossé, Y., Paré, G., & Meyre, D. (2019). Benefits and limitations of genome-wide association studies. Nature Reviews Genetics, 20(8), 467–484. 10.1038/s41576-019-0127-1

Taylor, S. A., Larson, E. L., & Harrison, R. G. (2015). Hybrid zones: windows on climate change. Trends in Ecology & Evolution, 30(7), 398–406. 10.1016/j.tree.2015.04.010

van Esse, H. P., Reuber, T. L., & van der Does, D. (2020). Genetic modification to improve disease resistance in crops. New Phytologist, 225(1), 70–86. 10.1111/nph.15967

Wang, L., Chen, S., Peng, A., Xie, Z., He, Y., & Zou, X. (2019). CRISPR/Cas9-mediated editing of CsWRKY22 reduces susceptibility to Xanthomonas citri subsp. citri in Wanjincheng orange (Citrus sinensis (L.) Osbeck). Plant Biotechnology Reports, 13(5), 501–510. 10.1007/s11816-019-00556-x

Wegrzyn, J. L., Liechty, J. D., Stevens, K. A., Wu, L. S., Loopstra, C. A., Vasquez-Gross, H. A., Neale, D. B. (2014). Unique features of the loblolly pine (Pinus taeda L.) megagenome revealed through sequence annotation. Genetics, 196(3), 891–909. 10.1534/genetics.113.159996

Wei, T. & Simko, V. (2017). R package “corrplot”: Visulaization of a correlation matrix. https://github.com/taiyun/corrplot

Weiss, M., Sniezko, R. A., Puiu, D., Crepeau, M. W., Stevens, K., Salzberg, S. L., De La Torre, A. R. (2020). Genomic basis of white pine blister rust quantitative disease resistance and its relationship with qualitative resistance. The Plant Journal, 104(2), 365–376. 10.1111/tpj.14928

Welsh, C., Lewis, K., & Woods, A. (2009). The outbreak history of Dothistroma needle blight: an emerging forest disease in northwestern British Columbia, Canada. Canadian Journal of Forest Research, 39(12), 2505–2519. 10.1139/X09-159

Wheeler, N. C. & Guries, R. P. (1987). A quantitative measure of introgression between lodgepole and jack pines. Canadian Journal of Botany, 65, 1876–1885. 10.1139/b87-257

White, T. L. (2004). TREE BREEDING, PRINCIPLES | Breeding Theory and Genetic Testing. In J. Burley (Ed.), Encyclopedia of Forest Sciences (pp 1551–1561). University of Oxford, Oxford, UK: Elsevier.

Wilson, J. F. & Goldstein, D. B. (2000). Consistent long-range linkage disequilibrium generated by admixture in a Bantu-Semitic hybrid population. The American Journal of Human Genetics, 67(4), 926–935. 10.1086/303083

Wood, L. J., Hartley, I. D., & Watson, P. (2009). Determining hybridization in jack pine and lodgepole pine from British Columbia. Wood and Fiber Science, 41, 386–395.

Woods, A. J. (2003). Species diversity and forest health in northwest British Columbia. The Forestry Chronicle, 79(5), 892–897. 10.5558/tfc79892-5

Woods, A. J., Coates, K. D., & Hamann, A. (2005). Is an unprecedented Dothistroma needle blight epidemic related to climate change? BioScience, 55(9), 761–769. 10.1641/0006-3568(2005)055[0761:IAUDNB]2.0.CO;2

Wu, M. C., Kraft, P., Epstein, M. P., Taylor, D. M., Chanock, S. J., Hunter, D. J., & Lin, X. (2010). Powerful SNP-set analysis for case-control genome-wide association studies. American Journal of Human Genetics, 86(6), 929–942. 10.1016/j.ajhg.2010.05.002

Wu, T. D. & Watanabe, C. K. (2005). GMAP: a genomic mapping and alignment program for mRNA and EST sequences. Bioinformatics, 21(9), 1859–1875. 10.1093/bioinformatics/bti310

Yeaman, S., Hodgins, K. A., Lotterhos, K. E., Suren, H., Nadeau, S., Degner, J. C., Aitken, S. N. (2016). Convergent local adaptation to climate in distantly related conifers. Science, 353(6306), 1431–1433. 10.1126/science.aaf7812

Yeatman, C. W. & Teich, A. H. (1969). Genetics and breeding of jack and lodgepole pines in Canada. The Forestry Chronicle, 45(6), 428–433. 10.5558/tfc45428-6

Zhou, H., Bai, S., Wang, N., Sun, X., Zhang, Y., Zhu, J., & Dong, C. (2020). CRISPR/Cas9-mediated mutagenesis of MdCNGC2 in apple callus and VIGS-mediated silencing of MdCNGC2 in fruits Improve resistance to Botryosphaeria dothidea. frontiers in Plant Science, 11, 575477. 10.3389/fpls.2020.575477

