## Supplementary material for "Genetic architecture underlying response to the fungal pathogen *Dothistroma septosporum* in lodgepole pine, jack pine, and their hybrids": Figure S1-S4

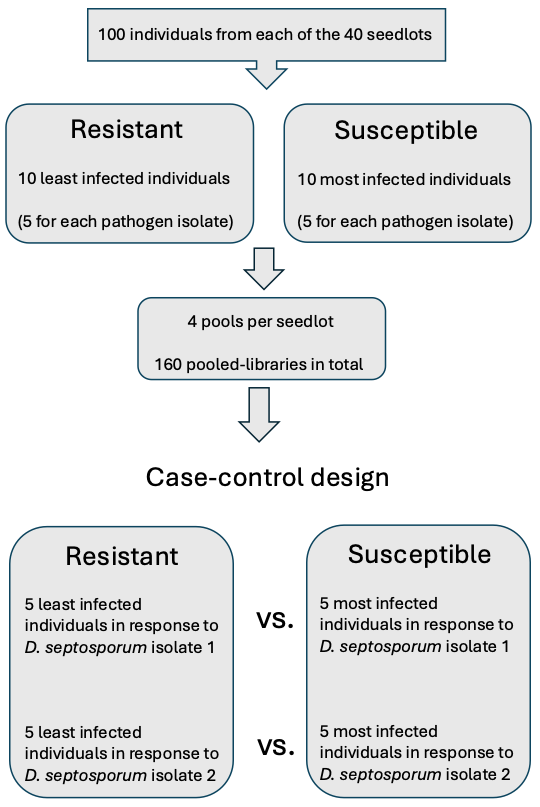

Figure S1. Flow chart of experimental procedure.

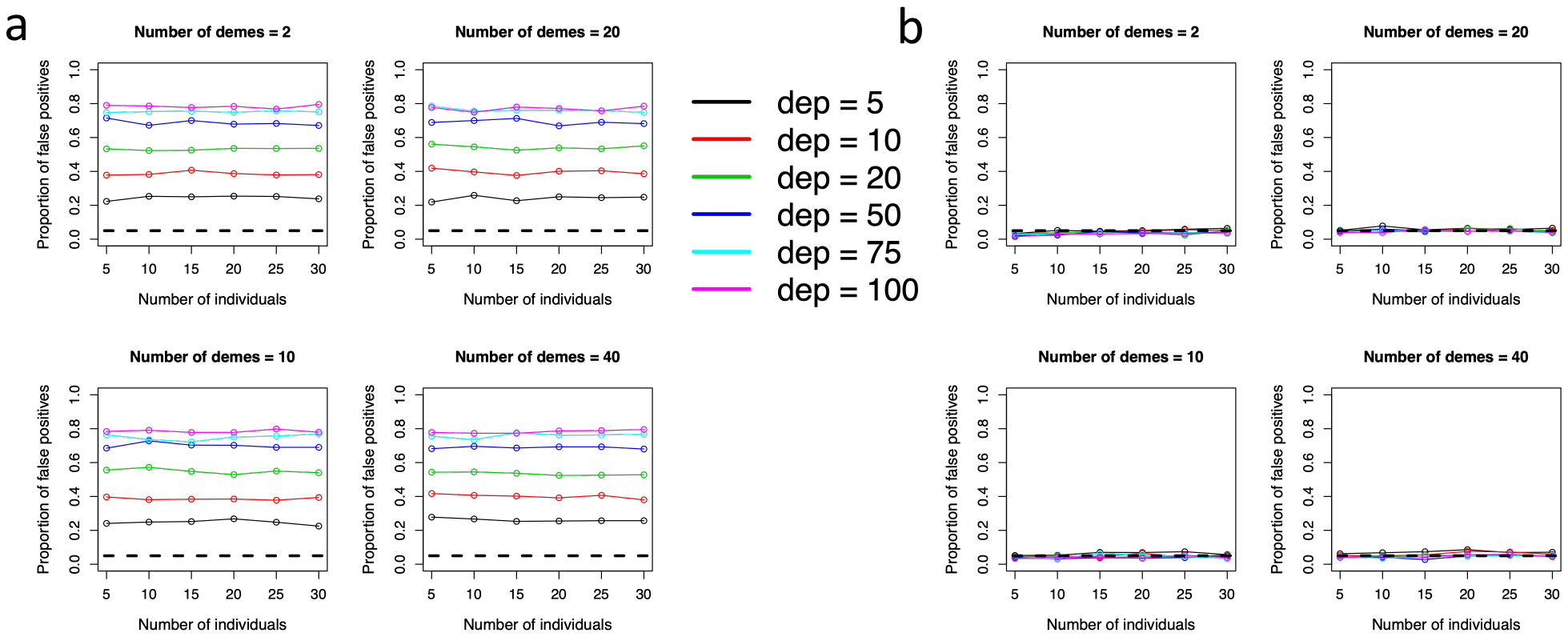

Figure S2. Power test of the CMH test. Simulation was run using uncorrected (a) and corrected allele counts (b) for pool-seq data. The corrected read counts were calculated by converting uncorrected allele counts to allele frequencies and then multiplying the ploidy (N=10) in each pool to acquire an approximate count that represents the real replication level. X-axis represents different numbers of individuals in each pool. dep ⎯ number of reads sampled per individual.

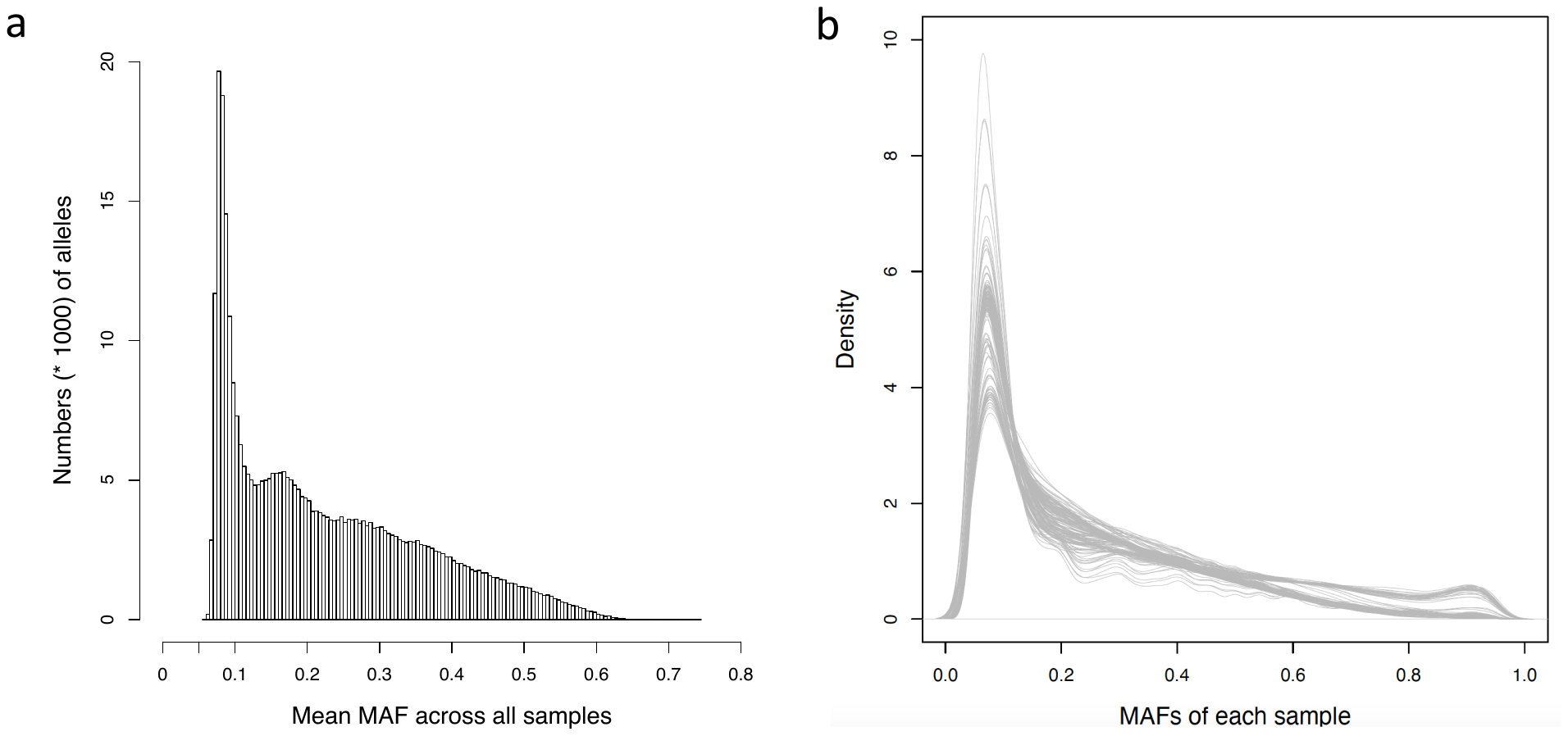

Figure S3. Minor allele frequency (MAF) for filtered SNPs across 160 pooled samples. a) Distribution of mean MAF across all samples. b) Density of MAFs of each sample. Each line represents a pooled sample.

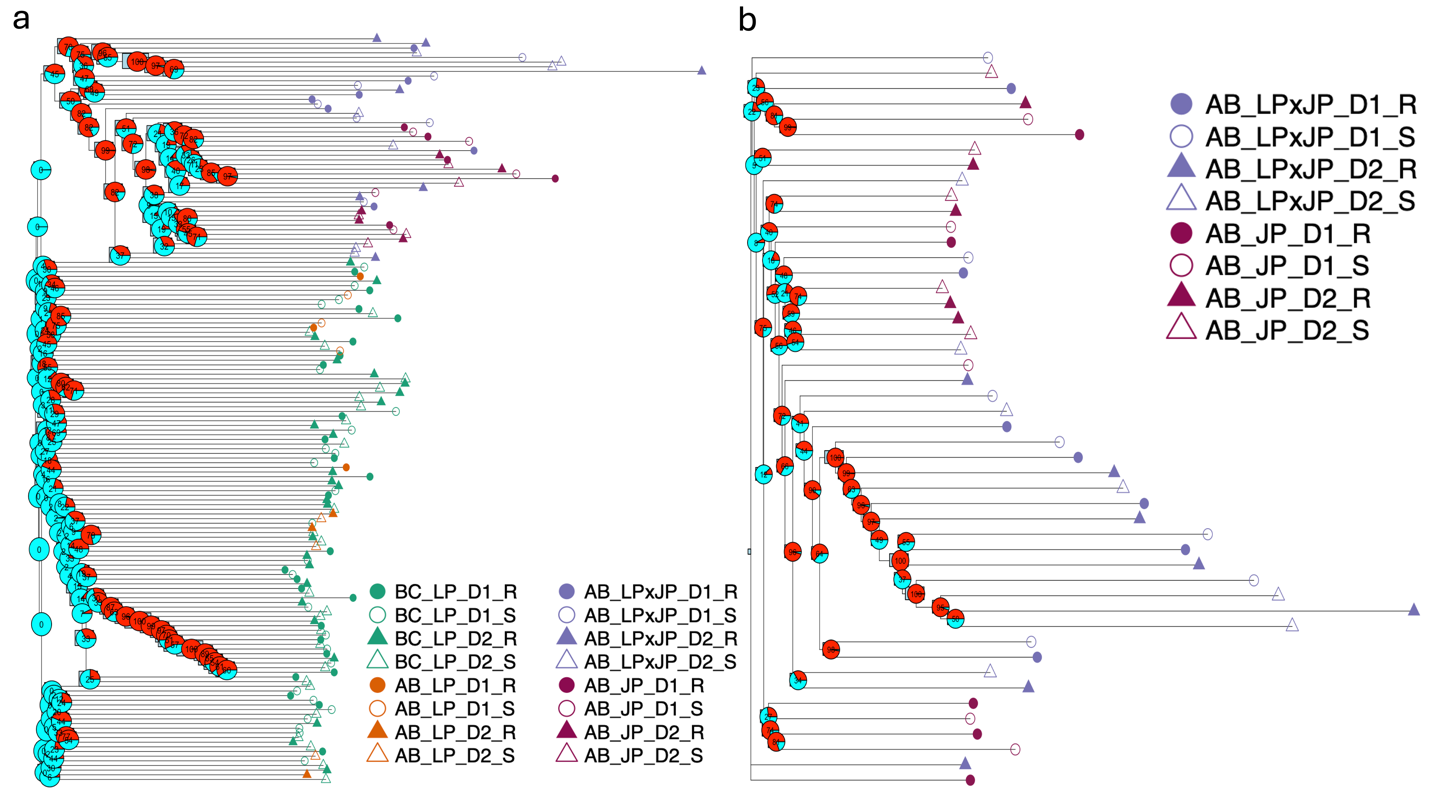

Figure S4. Phylogenetic tree constructed using the Neighbour-Joining algorithm for the 160 pooled samples (a) and for the 48 hybrid and jack pine samples (b). The bootstrap analysis was performed using 5000 bootstrap replicates. The bootstrap values are displayed as node labels. The red proportion in the pie represents the support percentage; the cyan proportion in the pie represents the non-support percentage. BC_LP ⎯ *Pinus contorta* (lodgepole pine) samples from British Columbia; AB_LP ⎯ *Pinus contorta* (lodgepole pine) samples from Alberta; AB_LPxJP ⎯ *Pinus contorta* × *P. banksiana* hybrid samples from Alberta; AB_JP ⎯ *Pinus banksiana* (jack pine) samples from Alberta. D1 ⎯ *D. septosporum* isolate 1; D2 ⎯ *D. septosporum* isolate 2. R ⎯ resistant tree; S ⎯ susceptible tree.
